# SARS-CoV-2 ORF3a Protein Impairs Syncytiotrophoblast Maturation, Alters ZO-1 Localization, and Shifts Autophagic Pathways in Trophoblast Cells and 3D Organoids

**DOI:** 10.1101/2024.09.25.614931

**Authors:** Deepak Kumar, Rowan M. Karvas, Brittany R. Jones, Eliza R. McColl, Emily Diveley, Sukanta Jash, Surendra Sharma, Jeannie C. Kelly, Thorold W. Theunissen, Indira U. Mysorekar

## Abstract

SARS-CoV-2 infection poses a significant risk to placental physiology, but its impact on placental homeostasis is not well understood. We and others have previously shown that SARS-CoV-2 can colonize maternal and fetal placental cells, yet the specific mechanisms remain unclear. In this study, we investigate ORF3a, a key accessory protein of SARS-CoV-2 that exhibits continuous mutations. Our findings reveal that ORF3a is present in placental tissue from pregnant women infected with SARS-CoV-2 and disrupts autophagic flux in placental cell lines and 3D stem-cell-derived trophoblast organoids (SC-TOs), impairing syncytiotrophoblast differentiation and trophoblast invasion. This disruption leads to protein aggregation in cytotrophoblasts (CTB) and activates secretory autophagy, increasing CD63+ extracellular vesicle secretion, along with ORF3a itself. ORF3a also compromises CTB barrier integrity by disrupting tight junctions via interaction with ZO-1, mediated by its PDZ-binding motif, SVPL. Colocalization of ORF3a and ZO-1 in SARS-CoV-2-infected human placental tissue supports our *in vitro* findings. Deleting the PDZ binding motif in the ORF3a protein (ORF3a-noPBM mutant) restored proper ZO-1 localization at the cell junctions in an autophagy-independent manner. Lastly, we demonstrate that constitutive ORF3a expression induces SC-TOs to transition towards a secretory autophagy pathway likely via the PBM motif, as the ORF3a-NoPBM mutants showed a significant lack of CD63 expression. This study demonstrates the functional impact of ORF3a on placental autophagy and reveals a new mechanism for the activation of secretory autophagy, which may lead to increased extracellular vesicle secretion. These findings provide a foundation for exploring therapeutic approaches targeting ORF3a, specifically focusing on its PBM region to block its interactions with host cellular proteins and limiting placental impact.

## Introduction

The emergence of severe acute respiratory syndrome coronavirus 2 (SARS-CoV-2) in 2019 led to COVID-19, a global pandemic that killed at least 6 million people by 2023^1,2^. Much is still unknown about disease pathogenesis and short- and long-term effects of COVID-19, particularly the effects of infection on pregnant women and their fetuses^3,4^. Although transmission from mother to fetus is rare, maternal infection has been associated with placental pathology, fetal/neonatal neurodevelopmental changes^5–7^, and increased risk of pregnancy-related disorders^8–12^, including a strong association with preeclampsia^6,12^. Placental pathologies and adverse pregnancy outcomes persist even after infections during pregnancy resolve^13,14^, indicating that the effects of SARS-CoV-2 infection can be long-lasting. Recently, long COVID has been identified as a novel multi-organ condition^15,16^, and can potentially impact women post-partum. Although vaccinations have reduced the overall burden of infections and are considered safe during pregnancy, rates of vaccination remain low amongst pregnant individuals^17^ ^10,18–21^. Thus, there is still a great need to understand how SARS-CoV-2 affects pregnancy and causes both short- and long-term sequelae.

The placenta serves as a physical and immunological barrier between the mother and the fetus^22,23^. Although incidence of SARS-CoV-2 vertical transmission is rare, cells composing the placenta express the SARS-CoV-2 entry receptor ACE2 and we and others have observed SARS-CoV-2 proteins and viral RNA in different compartments of the placenta over the course of gestation^24,25^. This demonstrates that SARS-CoV-2 can successfully invade the placenta and could have lasting effects on placental biology and function. However, the exact mechanisms through which SARS-CoV-2 impacts placental function remain unclear.

Autophagy, a crucial physiological process, plays a vital role in placental development by maintaining cellular homeostasis, and its dysregulation is associated with pregnancy complications such as preterm birth, miscarriage, and growth restrictions^26,27^. Moreover, autophagy is thought to play a pivotal role in preserving epithelial barrier function by regulating the transport and degradation of tight junction proteins (TJPs)^28,29^. Thus, autophagy is required for proper placental function and barrier formation. Changes in the expression of autophagy-related genes, disturbance in the regulation of autophagic proteins, and the disruption of autophagy signaling pathways have been reported in lung tissue and immune cells of COVID-19 patients^30–32^. Furthermore, autophagy is typically a degradative process where cellular components are enclosed in autophagosomes and sent to lysosomes for breakdown. However, when autophagosome-lysosome fusion is blocked, secretory autophagy process is initiated releasing undigested materials via extracellular vesicles^33,34,35–37^. Whether and how these processes impact placental trophoblast homeostasis is not known.

The SARS-CoV-2 accessory protein ORF3a has been shown to disrupt autophagy by blocking fusion of autophagosomes and lysosomes^38,39^. ORF3a has become a significant pathogenic element in the pathophysiology of SARS-CoV-2 and, similar to Spike protein, exhibits considerable mutability^38,40^; approximately175 mutations in the ORF3a protein have been identified across different SARS-CoV-2 variants^41^. These alterations are associated with a newly identified function of ORF3a, specifically its capacity to inhibit autophagosome-lysosome fusion, a characteristic not seen in ORF3a protein of SARS-CoV^42^. This suggests that placental invasion by SARS-CoV-2 could potentially disrupt placental autophagy, leading to altered placental function and ultimately, adverse pregnancy outcomes. Indeed, previous studies have established that disruption in autophagic flux is implicated in adverse pregnancy outcomes such as preeclampsia^26^, which is strongly associated with COVID-19^12^.

Here, we show that SARS-CoV-2 ORF3a is present in placental tissue from infected pregnant women and mechanistically show that it disrupts autophagic flux in placental cell lines and stem cell-derived 3D organoid models (SC-TOs)^43^. This disruption impairs syncytiotrophoblast differentiation, extravillous trophoblast invasion, leads to aberrant protein aggregation, activates secretory autophagy, and increases extracellular vesicle secretion, including secretion of ORF3a itself. ORF3a also compromises cytotrophoblast barrier integrity by interacting with ZO-1 via its PDZ-binding motif (PBM). Deleting this motif restores ZO-1 localization and prevents secretory autophagy. Together, our findings uncovered a novel molecular and cellular mechanism through which SARS-CoV-2 ORF3a disrupts placental function and compromises placental syncytial integrity.

## Results

### ORF3a impairs autophagic flux in placental trophoblasts

Immunostaining of placental tissue from uninfected and SARS-CoV-2-infected women reveals that ORF3a is present in SARS-CoV-2-positive placentas including in syncytiotrophoblasts (STBs) (**Supp. Fig. 1A-B**), indicating viral colonization and replication in the placenta, even if the mother is not actively infected at the time of tissue collection, consistent with previous reports^24,44,45^. We sought to determine the impact of SARS-CoV-2 ORF3a on placental trophoblast autophagy. JEG-3 cells^46^, a model of cytotrophoblasts (CTBs), were transfected with plasmids encoding the SARS-CoV-2 proteins ORF3a, ORF3b, Nucleocapsid (N), NSP6 and vector backbone as a control. After 24 hours, only ORF3a-transfected cells expressed significantly higher levels of the autophagy markers LC3B (**Fig. 1A-B**) and P62 (**Fig. 1C-D**), indicative of blocked autophagic flux. Similarly, forskolin-treated syncytialized JEG-3 cells, a model of STBs marked by increased HCG-β expression (**Fig. 1E**), showed increased expression of LC3B and P62 (**Fig. 1F and 1G**). Together, these results indicate that ORF3a specifically blocks autophagic flux in both CTBs and STBs. Consistent with this, we observed increased P62 and LC3 staining in SARS-CoV-2-infected placentas (**Sup Fig.1C-D**). As autophagy is involved in CTB-to-STB differentiation^47^, we examined the impact of ORF3a on HCG-β expression in syncytialized JEG-3. STBs transfected with ORF3a exhibited decreased HCG-β expression (**Fig. 1H**), suggesting that ORF3a disrupts the differentiation of CTBs into STBs.

**Figure 1:**
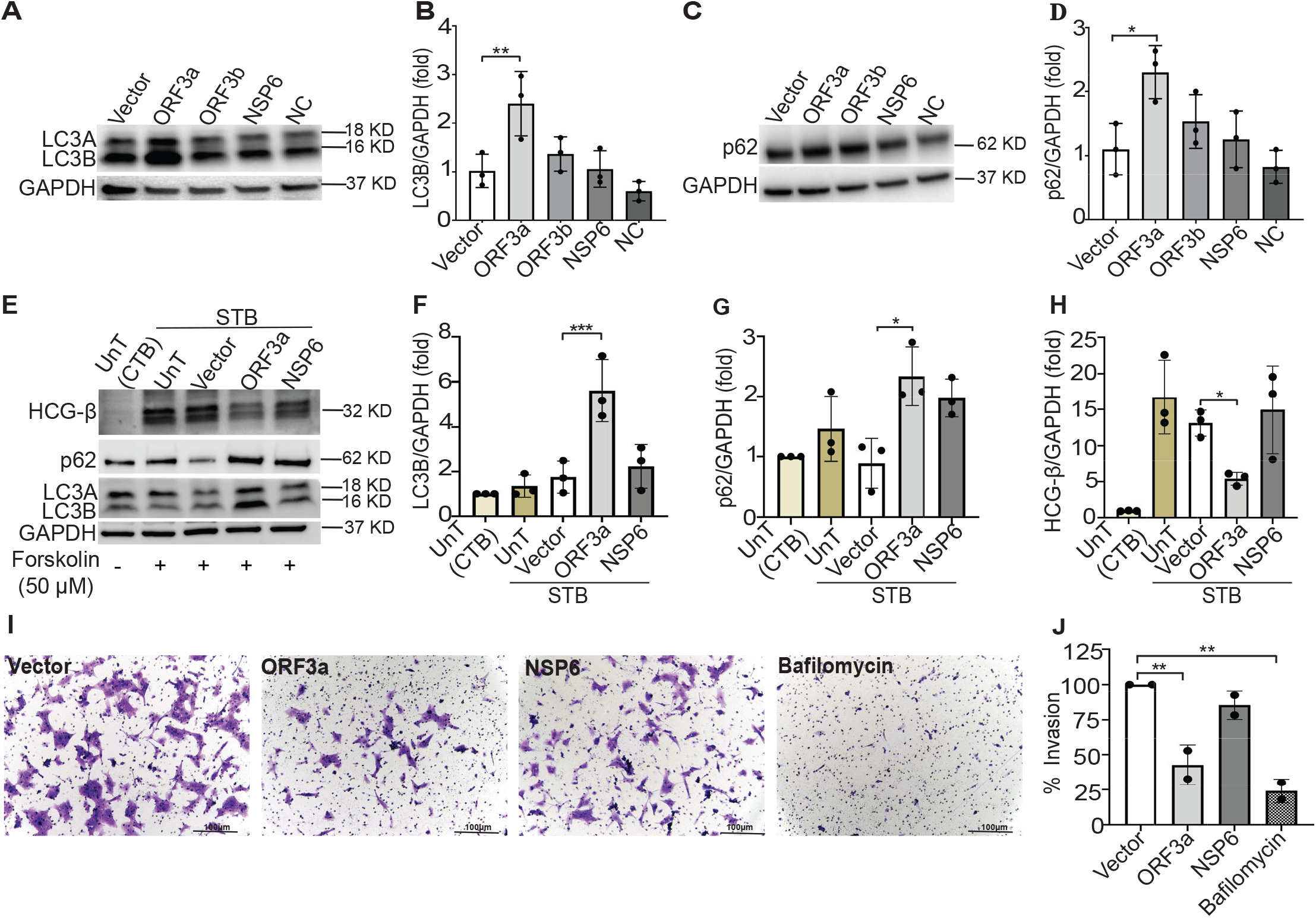
ORF3a blocks autophagy flux in CTBs, STBs and reduces Invasion in EVTs. **(A)** Jeg-3 cells transfected with SARS-CoV-2-encoded plasmids for 24 h, followed by Western blot for LC3B protein expression. **(B)** Densitometric quantification of LC3B protein expression normalized to GAPDH as loading control using Image Lab software (Data of three independent experiments were represented as mean ± SD, *p :S 0.05; ANOVA comparison test). **(C)** p62 protein expression detection by Western blot from Jeg-3 transfected cells, **(D)** followed by densitometric quantification of data acquired from three independent experiments (mean ± SD, *p :S 0.05; ANOVA comparison test). **(E)** A representative Western blot of forskolin (50µM) treated Jeg-3 cells as STB models which are also transfected with SARS-CoV-2 plasmids for 24 h and analyzed for expression of LC3B,p62 and HCG-B. Expression of LC3B **(F)** and P62 **(G)** increases significantly in ORF3a transfected STBs whereas HCG-B **(H)** shows reduced expression (Data of three independent experiments were represented as mean ± SD, *p :S 0.05 and ***p :S 0.001). **(I)** Brightfield microscopy after crystal violet staining of HTR-8 cells transfected with SARS-CoV-2 plasmids, also treated with bafilomycin for Matrigel invasion assay. **(J)** The quantification invasion was represented as percentage calculated after counting number of cells that invaded the Matrigel membrane from two independent experiments, analyzing a total of 10 regions of interest (ROIs). (Data presented as mean ± SD, **p :S 0.01)

Extravillous trophoblasts (EVTs) differentiate from CTBs and perform the essential function of invading maternal decidua to remodel spiral arteries^27,48^. The autophagy pathway has been shown to play a crucial role in trophoblast invasion function^48^; thus, we next examined whether SARS-COV-2 ORF3a interferes with this process using HTR-8/SVneo cells, a widely used *in vitro* model representing EVTs^49^.

Transfected cells were seeded onto Matrigel-coated trans-well inserts to quantify their invasive potential. ORF3a-transfected EVTs exhibited a significant decrease in their ability to invade through the Matrigel matrix compared to vector controls, suggesting a disruption of cell invasion mechanisms (**Fig. 1I and 1J**). Similar results were obtained when EVT cells were treated with bafilomycin A1 to model blocked autophagic flux, suggesting that the reduced invasion observed in ORF3a-transfected cells likely occurs because of blocked autophagic flux. Together, these results demonstrate that SARS-CoV-2 ORF3a blocks autophagic flux in multiple trophoblast cell types and impairs trophoblast differentiation and invasion.

### ORF3a induces protein aggregation

Canonical autophagy is typically degradative, clearing misfolded aggregated protein complexes^50^. Defects in this process have been demonstrated to induce the accumulation of aggregated proteins in the placenta, which is further associated with deleterious pregnancy outcomes such as pre-eclampsia^26,51^. Thus, we next sought to determine whether ORF3a leads to protein aggregation in the placenta. Confocal imaging of JEG-3 cells transfected with ORF3a revealed accumulation of Proteostat dye, which binds specifically to aggregated proteins (**Fig. 2A** and **2C**). Furthermore, Proteostat-positive aggregates colocalized with P62 in ORF3a-transfected JEG-3 (**Fig**. **2A** and **2D**), suggesting that ORF3a-mediated blockage of autophagy prevents the degradation of P62-marked proteins, leading to their accumulation. Indeed, cells exposed to bafilomycin A1 exhibited a significant increase in Proteostat staining (**Fig. 2B** and **2E**), supporting the hypothesis that the accumulation of protein aggregates occurs due to disrupted autophagic degradation. Consistent with the *in vitro* results, human placentas from SARS-CoV-2-infected pregnancies exhibited increased Proteostat staining (**Sup Fig. 1E**). Together, this suggests that SARS-CoV-2 induces protein aggregation through ORF3a-mediated blockage of autophagic flux.

**Figure 2:**
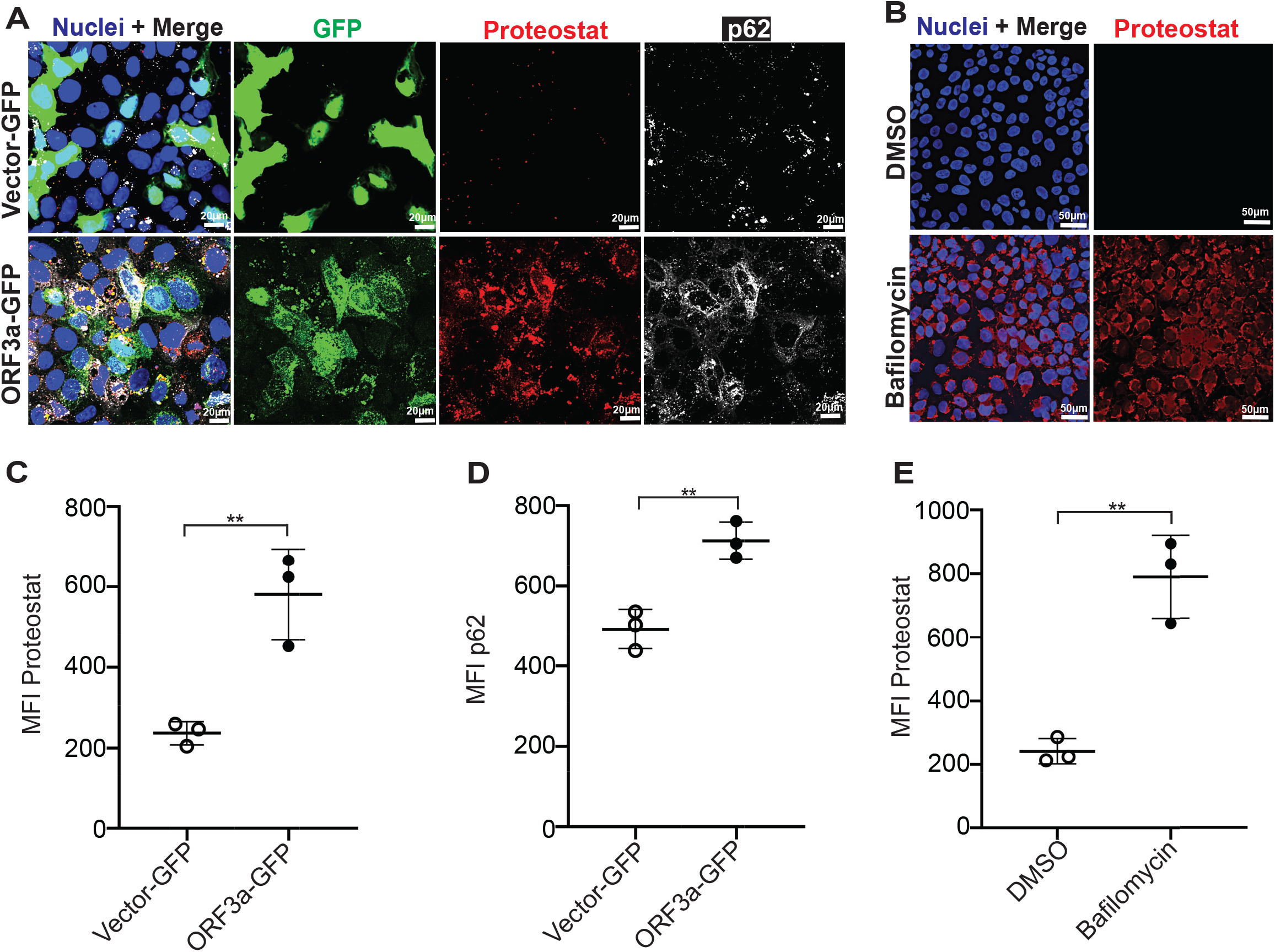
ORF3a induces protein aggregation. **(A)** Representative confocal images of Jeg-3 CTBs transfected with ORF3a-GFP (green) exhibit enhanced proteostat staining (red) and increased p62 staining (white) relative to Vector-GFP transfected CTBs. **(B)** Confocal imaging of Jeg-3 cells treated with bafilomycin and DMSO (vehicle control) and stained with proteostat dye (red). **(C)** Data quantification from three independent experiments for panel (A) after averaging mean fluorescence intensity (MFI) from a minimum of six images per condition and per biological replicate and shown as mean ± SD, **p < 0.05, using unpaired two-tailed Student’s *t* test. **(D)** MFI of P62 was quantified from three independent experiments after averaging four images per condition and per biological replicate and observed colocalization with proteostat and ORF3a-GFP (mean ± SD, unpaired two-tailed Student’s t test). Scale bar=20 µm. **(E)** Mean fluorescence intensity (MFI) quantification for panel (B) shows increased proteostat staining in bafilomycin treated Jeg-3 cells, data of three independent experiments with averaging MFI from minimum of four images per condition per replicate, mean ± SD, **p < 0.05, unpaired two-tailed Student’s *t* test. Scale bar=50 µm.

### ORF3a-mediated disruption of canonical autophagy leads to activation of secretory autophagy and induces upregulation of extracellular vesicle secretion

In the absence of autophagosome-lysosome fusion or malfunctioning lysosomes, autophagy switches to secretory autophagy, which is otherwise a degradative mechanism^36,52^. When in this state, autophagosomes or malfunctioning lysosomes approach the plasma membrane and release their cargo into the extracellular space. Thus, given that ORF3a appears to block canonical autophagic flux, we investigated the possibility that secretory autophagy could be triggered in the placenta by ORF3a. We first investigated whether ORF3a transfection in trophoblast cells increases CD63+ vesicle formation, as CD63 identifies EVs produced during secretory autophagy^53,54^. Indeed, ORF3a-transfected JEG-3 cells had significantly more CD63+ vesicles compared to vector control, indicating a shift towards an active secretory pathway (**Fig. 3A** and **3B**). Furthermore, CD36+ vesicles colocalized with ORF3a (**Fig. 3A**), suggesting that ORF3a may be secreted through these EVs during secretory autophagy. To test this, conditioned culture media from ORF3a-transfected JEG-3 cells was harvested and cleared of any cellular debris before being added to new, non-transfected JEG-3 cells. Confocal microscopy showed that JEG-3 cells treated with the conditioned media exhibited high levels of ORF3a protein (**Fig. 3C**), suggesting that ORF3a had been secreted into the conditioned media. Moreover, cells treated with the conditioned media exhibited colocalization of ORF3a with Lamp1-positive lysosomes (**Fig. 3D**), suggesting endocytosis of ORF3a from the conditioned media. Taken together, these results suggest the possibility that when canonical autophagy is blocked due to ORF3a expression, secretory autophagy occurs, resulting in ORF3a secretion, exosome packaging, and endocytosis into neighboring cells, even in the absence of active viral replication.

**Figure 3.**
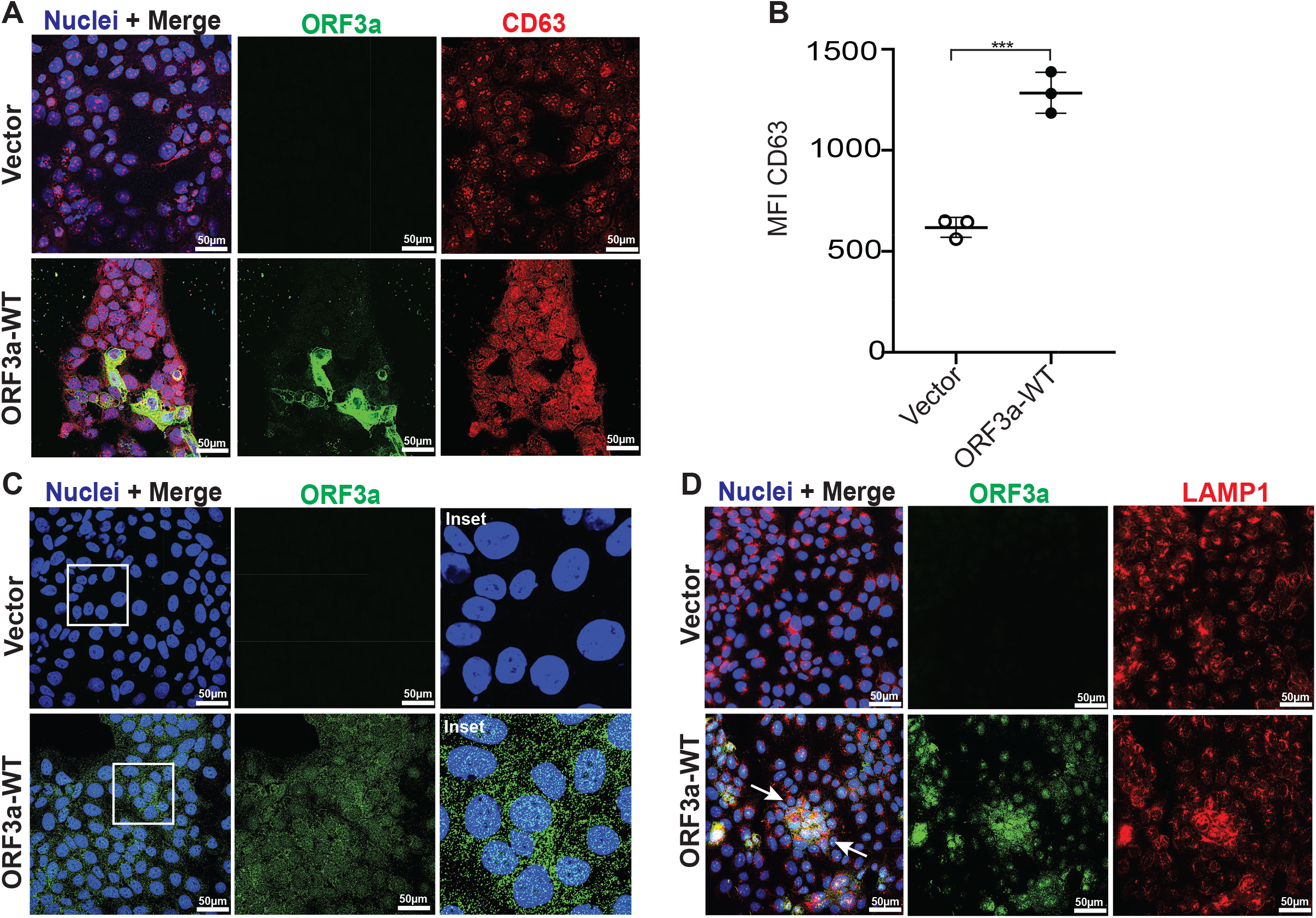
ORF3a induces secretory autophagy and is secreted with EVs. **(A)** Confocal representative IF images from three independent experiments show enhanced formation of CD63+ (green) vesicles in ORF3a-WT transfected (red) Jeg-3 cells as compared to vector control (Scale bar = 50 μm). **(B)** Comparison of CD63+ vesicles MFI between vector and ORF3a-WT transfected cells with significant differences (mean ± SD, ***p<0.05) determined by unpaired two-tailed Student’s t test, based on 4 images per condition per group from three independent experiments. **(C)** Confocal IF images from three independent experiments show Jeg-3 cells treated with conditioned media from Vector and ORF3a-WT (green) transfected cells, with nuclei stained using DAPI (Scale bar = 50 μm). **(D)** Jeg-3 cells exposed to ORF3a-containing (green) conditioned media demonstrate endocytosis and colocalization (yellow) with Lamp1 (red) in healthy cells (Scale bar = 50 μm).

### ORF3a interacts with ZO-1 via the PBM motif and alter its localization in trophoblasts

Canonical autophagy plays a role in preserving the structural integrity of epithelial cell-cell junctions by controlling the trafficking and recycling of the ZO-1 protein^36^. However, given our findings that ORF3a induces a transition from conventional autophagy to secretory autophagy, we sought to determine the influence of ORF3a on the barrier integrity of placental trophoblasts. At its C-terminus cytosolic regions, ORF3a harbors a short linear motif (SLiM) designated PBM (PDZ binding motif) (amino acid sequence *SVPL*) (**Fig. 4A**), known to interact with cellular PDZ domain-containing proteins which are typically involved in cell junction and polarity^55^. Thus, we hypothesized that ORF3a may be capable of directly binding to the PDZ domain of ZO-1. However, the commercially available pLVX-EF1alpha-ORF3a-2xStrep plasmid has a 2xStrep tag on the C-terminus of ORF3a, which would inhibit the direct binding capacity of its PBM (**Fig. 4A**). Thus, we used restriction enzyme-based cloning to eliminate the 2x-Strep tag by inserting a stop codon upstream of the tag sequence and added EcoRI and BamHI sites. Moreover, we altered the C-terminal PBM motif of ORF3a by introducing a stop codon immediately downstream of the PBM site and incorporated a BamHI site post the new stop codon. The forward primer was designed to include an EcoRI site. Post-PCR, the amplicon was subjected to double digestion with EcoRI and BamHI, followed by ligation into the pLVX-EF1alpha vector. Successful cloning was validated through colony PCR, double digestion and Sanger sequencing (**Fig. 4B**).

**Figure 4:**
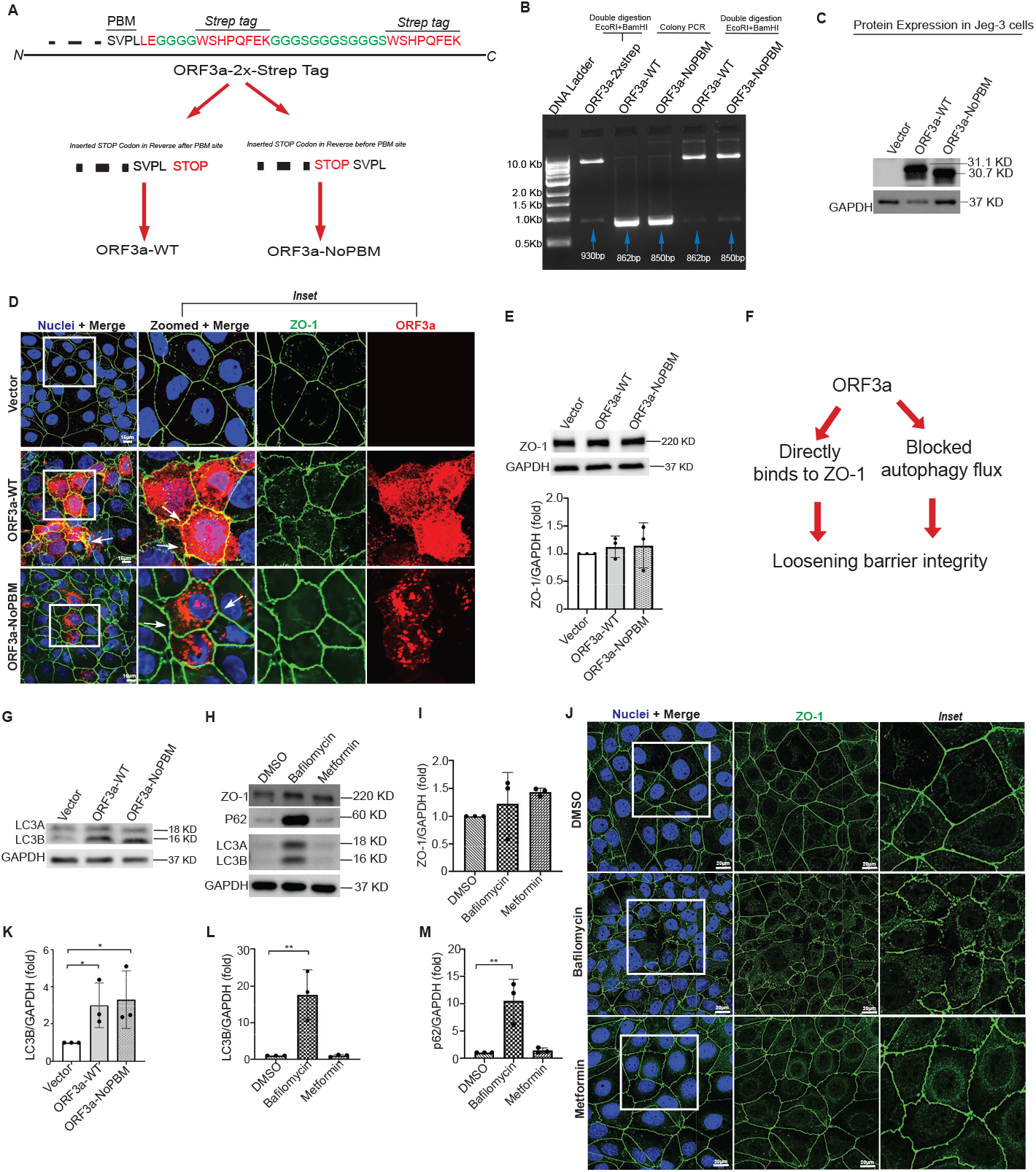
ORF3a directly associates with ZO-1 via PBM motif and alters its localization. (A) Schematic illustration of the constructed plasmids: ORF3a-WT (wild type or untagged) and ORF3a-NoPBM (with PBM motif deletion). **(B)** Clone confirmation by DNA gel electrophoresis demonstrates that colony PCR from positive colonies successfully integrated ORF3aWT (862 bp) and ORF3a-NoPBM (850 bp) genes at their respective sizes into the parent ORF3a-2X-Strep plasmid. This was further verified by double digestion, confirming the correct integration of genes with the pLVX backbone plasmid. **(C**) Western blot analysis of ORF3a-WT and ORF3a-NoPBM expression in transfected Jeg-3 cells. (D) Immunostaining of Jeg-3 cells transfected with vector control, ORF3a-WT and ORF3a-NoPBM shows co-localization of ORF3a-WT (in red) and ZO-1 (in green) as compared to ORF3a-NoPBM and Vector. Insets provide magnified views of the co-localization regions (Scale bar = 15µm). **(E)** Western blot of Jeg-3 cells transfected with ORF3a-WT and ORF3a-NoPBM show unchanged expression of ZO-1 as represented by data of three independent experiments with mean ± SD, p>0.05 as non-significant. **(F)** A schematic illustration of mechanism by which ORF3a impacting the ZO-1 localization. **(G)** LC3B expression analyzed through western blot shows increased trend in both ORF3a-WT and ORF3a-NoPBM. (Data presented as mean ± SD, *p :S 0.05) **(H)** Jeg-3 cells treated with bafilomycin, metformin and respective vehicle control DMSO, analyzed for expression levels of LC3B, p62 and ZO-1. (mean ± SD **p :S 0.01) **(I)** Quantification of ZO-1 expression for panel H shows unchanged level in both bafilomycin and metformin treatment. **(J)** Immunostaining and confocal imaging for Jeg-3 cells treated with DMSO, bafilomycin and metformin, representative images from three independent experiments show expression of ZO-1 (green) and nuclei stained with DAPI (Scale bar = 20µm). **(K)** Quantification of LC3B expression from panel G (mean ± SD, *p :S 0.05). **(L)** and **(M)** corresponds to the quantification of western blot for LC3B and p62 in bafilomycin and metformin treatment for panel H respectively (mean ± SD, **p < 0.05).

Newly generated plasmids ORF3a (termed ORF3a-WT) or ORF3a protein lacking SVPL (termed ORF3a-No-PBM) successfully expressed either form of ORF3a when transfected into JEG-3 cells (**Fig. 4C)**. When expressed in JEG-3 cells (**Fig. 4C**), ORF3a-WT co-localized strongly with ZO-1 (**Fig 4D; inset**). In contrast, ORF3a-No-PBM exhibited a near total lack of co-localization with ZO-1 (**Fig. 4D; inset**) Furthermore, while overall protein expression of ZO-1 was unchanged by either form of ORF3a (**Fig. 4E**), ORF3a-WT caused mis-localization of ZO-1 and breakage of tight junctions, while the PBM mutant did not (Fig. 4D; inset).This suggests that ORF3a directly binds and mis-localizes ZO-1, and that the PBM domain of ORF3a is required for this interaction. ORF3a-WT is distributed throughout the cell (**Fig. 4D; Inset**), whereas the ORF3a-NoPBM mutant is less widely distributed and only present in perinuclear regions. This also suggests that the PBM domain of ORF3a helps in intracellular transport of ORF3a. We further note colocalization of ORF3a and ZO-1 in SARS-CoV-2-infected human placenta samples (**Supp. Fig. 1B**).

We further observed that both ORF3a-WT and ORF3a-NoPBM mutant are capable of blocking autophagy flux as shown by an increase in LC3B expression (**Fig. 4G** and **4K**). Thus, it is possible that the impact of ORF3a on ZO-1 localization may occur indirectly through autophagy in addition to direct interaction between ORF3a and ZO-1 (**Fig. 4F**). Thus, to investigate the potential direct regulation of ZO-1 expression and localization by autophagy, JEG-3 cells were treated with either bafilomycin (autophagy inhibitor) or metformin (autophagy activator)^34^. Consistent with the impact of ORF3a, there was no overall change in the expression of ZO-1 in bafilomycin-treated cells as compared to controls (**Fig. 4H and 4I**). Bafilomycin treatment, which increased expression of LC3B and p62 indicative of blocked autophagic flux (**Fig. 4L and 4M**), mislocalized ZO-1 away from the cell membrane and scattered in the cytoplasm (**Fig. 4J**). In contrast, metformin treatment, in which LC3B and p62 levels were normalized due to proper autophagic flux, resulted in intact ZO-1 localization at the membrane. Overall, these results suggest that in addition to direct interaction between ORF3a and ZO-1, autophagy plays a role in appropriately localizing ZO-1 at the cell membrane.

### ORF3a reduces maturation and induces secretion of EVs in of 3D SC-TOs

To determine further whether the impact of ORF3a on autophagy and differentiation of CTBs to STBs also persists in 3D organoid models, we utilized human trophoblast stem cells (hTSCs)-derived 3D SC-TO models^43,56,57^. These 3D trophoblast organoids were created using hTSCs obtained from the first trimester of the human placenta^58^ that can differentiate into STBs and EVTs^43,56^. These models were successfully used to examine infections with Zika virus^43,59^ and SARS-CoV-2^43,60^, indicating their relevance in modelling placental vulnerability to pathogens and first trimester infections in placental cells.

To assess the effect of ORF3a on the three-dimensional development of SC-TOs, we initially established a CT-30 hTSC cell line that stably expresses ORF3a-WT, ORF3a-NoPBM and vector controls through lentiviral transduction. Subsequently, these cells were used to generate SC-TOs using our previously published protocol^43^. We documented the development of 3D SC-TOs through bright-field imaging on the 5th and 10th days to monitor morphological changes. Our findings indicate that SC-TOs derived from ORF3a-expressing hTSCs exhibit hindered development compared to the vector controls (**Fig. 5A**). Notably, the average diameter of ORF3a-expressing SC-TOs was significantly smaller than that of the vector control SC-TOs at both observed time points (**Fig. 5A and 5B**). The autophagic flux analysis showed increased expression of LC3b and p62 in ORF3a-expressing SC-TOs (**Fig. 5C-5E**) as compared to vector controls, consistent with our findings in placental cell lines.

**Figure 5:**
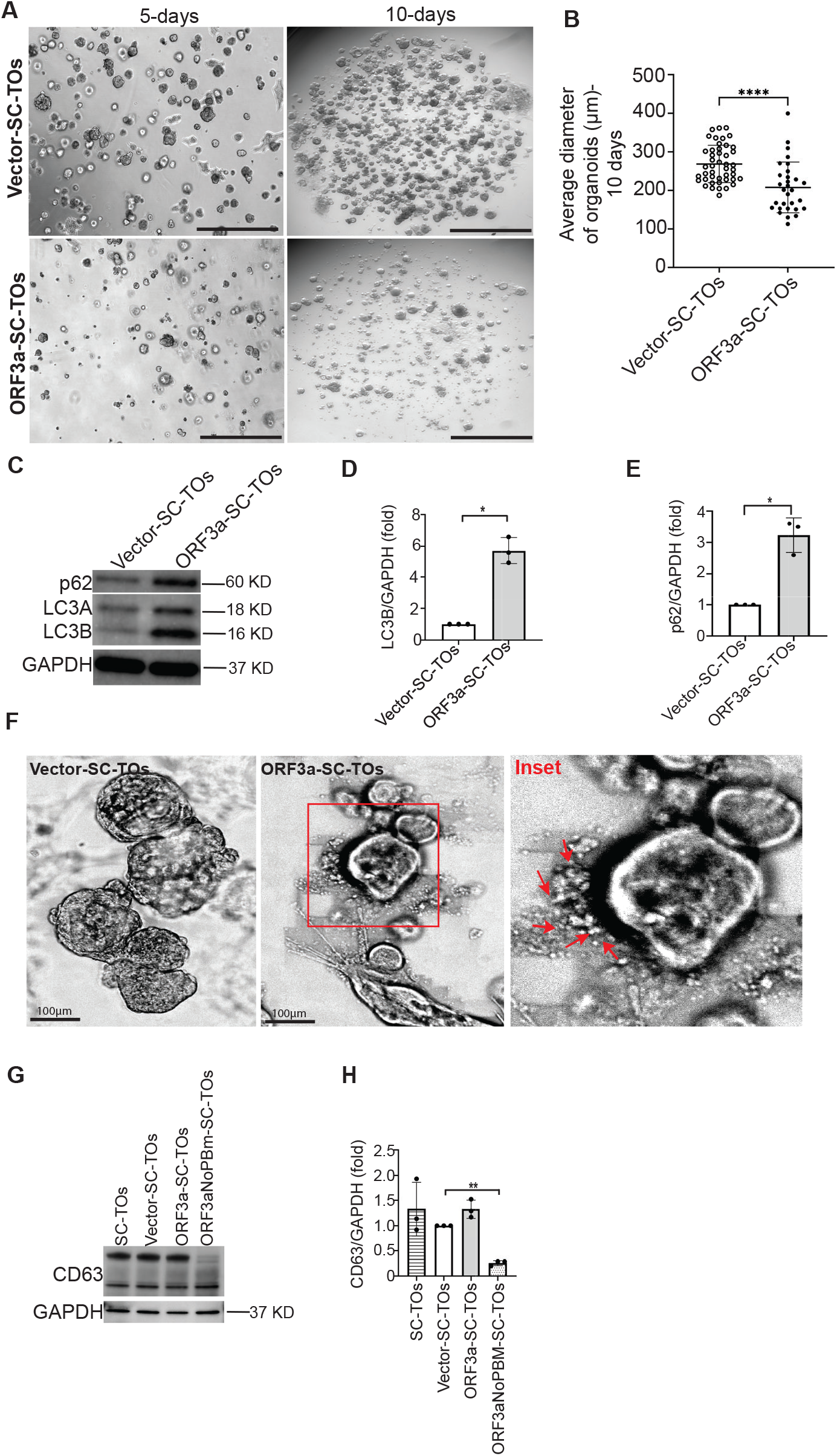
ORF3a reduced maturation and induces secretion of EVs in of 3D SC-TOs. **(A)** Bright field images captured at 5x magnification of SC-TOs developed from CT30-Vector and CT30-ORF3aWT, demonstrating growth at 5 and 10 days (Scale Bar=1000 μm). **(B)** Comparison of average organoid diameter between CT-30-Vector and CT-30-ORF3a-WT, with significant difference (mean ± SD, ****p<0.0001) determined by Mann-Whitney test, based on nearly 30 ROIs per group. **(C)** A representative Western blot of proteins isolated from vector-SC-TOs and ORF3a-WT-SC-TOs probed for LC3B and p62 expression. (D) and (E) represents the quantification of blot showing increased expression of LC3B and p62 in ORF3a-WT-SC-TOs (data represented from three independent experiments as mean ± SD, *p < 0.05) **(F)** Brightfield images of SC-TOs at 20x magnification illustrating the formation of extracellular vesicles (inset: red arrows) by CT-30-ORF3a-WT organoids (Scale Bar=100 μm). (G) Representative Western blot showing expression of CD63 where quantification **(H)** revealed reduced expression of CD63 in ORF3a-NoPBM-SC-TOs.

Moreover, we found that ORF3a-SC-TOs are encircled by numerous extracellular vesicles (EVs) (**Fig. 5F**), suggesting that ORF3a may induce these organoids to transition towards a secretory autophagy pathway. We next wondered whether these EVs were positive for CD63. Western blotting for cell lysates for CD63 revealed that expression of CD63 does not change in ORF3a-SC-TOs with respect to vector and wild type SC-TOs. CD63 is known to harbour a PBM^61^ and thus we wondered if CD63+ EV secretion was affected equally by ORF3a-WT and ORF3a-NoPBM. Interestingly, we found significantly reduced CD63 expression in ORF3a-NoPBM compared with ORF3a-WT-SC-TOs (**Fig. 5E and 5F**). These findings suggest that the PBM domain of ORF3a may interact with CD63and affect its secretion capacity.

## Discussion

This study demonstrates that SARS-CoV-2 ORF3a protein plays a key role in modulating autophagy and barrier function in trophoblasts, providing a potential mechanistic insight into placental dysfunction observed during COVID-19 infection. Our findings provide evidence that SARS-CoV-2, particularly through its ORF3a protein, significantly disrupts autophagic processes in placental trophoblasts, potentially impacting key aspects of placental function and integrity. The observed impairment of autophagic flux in cell lines as well as 3D SC-TO suggests a key role of autophagic processes in placental homeostasis. This disruption is further evidenced by the accumulation of Proteostat-positive aggregates in the Jeg-3 cells. In placentas affected by preeclampsia, a deficiency in autophagic processes has been consistently linked with the accumulation of protein aggregates, a pathological feature that contributes significantly to the severity of the preeclamptic phenotype^26,51^. These findings were further paralleled by the observations in lung tissue from COVID-19 patients, where an increase in both LC3B- and P62-positive cells was noted^32^. The elevation of LC3B and P62 mRNA levels in the lungs of these patients supports the hypothesis of an augmented autophagic initiation but also indicates a potential stopping of the autophagic process, resulting in protein accumulation^32^.

Furthermore, our study identified ORF3a as a significant disruptor of autophagic flux in placental trophoblasts. The increased levels of both LC3B and P62 in ORF3a-transfected cells, along with the pronounced increase in Proteostat staining in cells and human placental tissues, underscore the direct impact of ORF3a on autophagy and protein aggregation. Our findings are corroborated by existing studies that have identified the ORF3a protein as an inhibitor of autophagic flux^62,63^. Specifically, ORF3a has been demonstrated to interfere with critical steps in the autophagic pathway. Mechanistically, it has been shown that ORF3a can sequester VPS39, a component of the HOPS complex, which is essential for the fusion between autophagosomes and lysosomes^62,63^. This sequestration prevents the normal interaction between VPS39 and RAB7, a key regulator of membrane fusion events, resulting in a failure of autophagosome-lysosome fusion and subsequent impairment of the autophagic process. The reduction in trophoblast invasion and syncytialization upon ORF3a transfection or bafilomycin treatment mirrors the findings of impaired trophoblast function in cases of hypoxia or pharmacological inhibition of autophagy. This ties into the broader spectrum of placental pathologies, including pre-eclampsia, where aberrant autophagy has been repeatedly implicated^26,51,64^. The findings highlight a dual interference by ORF3a in STB function: impeding the completion of the autophagic process and potentially compromising STB maturation. These findings concur with prior research using a trophoblast stem cell model to examine the impact of SARS-CoV-2 infection on STB differentiation wherein a noticeable reduction in HCG-β secretion was observed, along with an incomplete differentiation profile in virus-infected cells^65^. Dysfunctional morphology and differentiation of syncytiotrophoblasts *in vivo* and impaired differentiation of deeply invasive extravillous trophoblasts^66^ are known contributors to the development of pre-eclampsia, a complication that manifests with increased frequency among pregnant women diagnosed with COVID-19^67,68^. Our work identifies a key target for potentially limiting the impact of SARS-CoV-2 infection in pregnant women.

Our study identifies molecular interactions between ORF3a and tight junction protein, Zona Occludens-1 (ZO-1). The altered distribution and expression levels of ZO-1 in ORF3a-transfected cells suggest that SARS-CoV-2 infection may compromise trophoblast barrier functionality, a hypothesis supported by the co-localization of ORF3a and ZO-1 in infected human term placentas. Our data also showed that autophagy inhibition by bafilomycin induced a mis-localization of ZO-1, a finding that is indicative of compromised junctional integrity in the presence of autophagy inhibition. Our findings that the disruption of ZO-1 localization is more pronounced in ORF3a-WT than in ORF3a-NoPBM-transfected cells hint at a direct interference mechanism that warrants further investigation. The fragmented pattern of ZO-1 suggests that ORF3a expression weakens the organized network of tight junctions essential for trophoblast barrier function by reducing autophagic flux and directly associating with ZO-1 protein. The visualization of ORF3a and its colocalization with ZO-1 in SARS-CoV-2-infected term placentas underscores the *in vivo* relevance of our *in vitro* findings and strengthens the proposed link between viral protein expression and compromised barrier integrity. This could explain the histopathological alterations^11,69–71^ observed in placentas from COVID-19 patients, aligning with other studies reporting similar findings. Also, our findings support prior work showing ZO-1 playing a role in CTB-STB differentiation^72^. ZO-1 is a critical component of the tight junction complex that regulates cell polarity and paracellular permeability, which are vital for the epithelial barrier function and structural integrity^73,74^. The colocalization of ORF3a and ZO-1 in SARS-CoV-2-infected placentas implicates the virus in potentially modulating the permeability of the placental barrier. This is particularly significant since the integrity of the STB layer is essential for the barrier against infections and for maintaining fetal-maternal exchange.

Our study further identified that ORF3a-induced impairment of canonical autophagy in trophoblasts could lead to activation of secretory autophagy and upregulation of CD63-associated EVs where ORF3a itself is secreted together with CD63. Our findings support prior observations that ORF3a can bind to Lamp1-positive lysosomes once it has been endocytosed^75^. This secretory phenotype was also consistently seen in our 3D SC-TOs stably expressing ORF3a-WT as compared to vector and wild type SC-TOs. Such a vesicle secretory phenotype has not been previously reported in any of the trophoblast organoid model under native conditons^43,76^, underscoring a driving role of ORF3a in inducing this secretory phenotype. Notably, the ORF3a-NoPBM mutant showed significant reduction of CD63 expression in 3D SC-TOs, indicating a possible role of the PBM motif in physical association with CD63. CD63 also has a PBM, which interacts with syntenin-1, promoting exosome formation and trafficking^77^. Our observation of lower expression of CD63 in cell lysates of 3D-SC-TOs expressing the ORF3a-noPBM mutant suggests the important involvement of ORF3a’s PBM domain in extracellular vesicle formation and its integration into CD63+ exosomes. When lysosomes become dysfunctional or autophagosomes are unable to fuse with lysosomes, secretory autophagy is activated to eliminate the damaged lysosomes, autophagosomes, or mitochondria^52^. Broadly, when intracellular trafficking halts, this pathway clears out cellular debris^78^. Also, this pathway has been associated with upregulation of EV formation and release in response to stress^35,37^. These EVs may contain proteins, lipids, and nucleic acids and have an impact on nearby or distant recipient cells^79^. Moreover, EVs have been noted to contain viral RNA during SARS-CoV-2 infection, indicating that they might aid in the propagation of the virus by endocytosis^80^. SARS-CoV-2 infection has been shown to enhance exosome secretion^80^ which may result in the generation of circulating exosomes that adversely affect the placenta and inhibit trophoblast differentiation and invasion^81^. Together, these investigations point to a novel method of ORF3a triggering secretory autophagy which enhances exosome secretion, and the physical interaction of ORF3a with exosomes, which may induce it to be secreted alongside cargo. These findings provide a foundation for exploring therapeutic approaches targeting ORF3a, specifically focusing on its PBM region to block its interactions with host cellular proteins and limiting placental impact.

In sum, our work provides new insights into potential viral mechanisms through which SARS-CoV-2 infection may exacerbate placental dysfunction and contribute to the clinical complications observed in pregnant women with COVID-19.

## Methods

Human samples were obtained from via random sampling from a pre-existing prospective cohort study of pregnant individuals during the COVID-19 pandemic enrolled at Barnes-Jewish Hospitals in St. Louis, MO from December 2021-July 2022 (Safety, Testing/Transmission, and Outcomes in Pregnancy with COVID-19 (STOP-COVID-19) study). Patients were serially evaluated for exposure to SARS-CoV-2 infection at enrollment. Antepartum infection was evaluated universal testing at delivery for SARS-CoV-2 using polymerase chain reaction (PCR) and/or antigen testing. This study was approved by the institutional review board (#202012075). The participants provided informed consent to sampling, storage and use of clinical samples. Placenta samples included villous tissue from COVID (+) and COVID (-) patients. The inclusion criteria included term births, singletons, unvaccinated against SARS-CoV-2. Exclusion criteria included pre-eclampsia, preterm or still births, and any other infections.

### Cell Culture

Human trophoblastic cell lines, including JEG-3 (ATCC HTB-36), and HTR-8/SVneo (ATCC CRL-3271), were propagated in Dulbecco’s Modified Eagle Medium/Nutrient Mixture F-12 (DMEM/F-12, GIBCO Cat. No. 11330032) enriched with 10% fetal bovine serum (FBS, Gibco, Cat. No. 16140071). The cultures were incubated in a controlled environment at 37 °C and an atmosphere containing 5% CO2. JEG-3 cells were induced to undergo syncytialization by treatment with 50μM Forskolin (Sigma, F6886) for a duration of 24 hours. For blocking and inducing autophagy pathways in cell lines, bafilomycin A1(Sigma, B1793) (100nM) and metformin (Sigma, 317240) (50μM) were used respectively.

### 3D SC-TO generation

3D SC-TOs were created using the CT30 hTSC 2D cell line, as previously described^43^. 2D hTSCs were grown in 6-well plates coated with Laminin521. Lamini521 was kept in wells and stored at 4°C overnight. Before employing these coated plates, they were incubated at 37°C for 1 hour. The medium was DMEM/F12 with 0.1 mM 2-mercaptoethanol, 0.2% FBS, 0.5% Penicillin-Streptomycin, 0.3% BSA, 1% ITS-X (Gibco, 51500), 1.5 μg/ml L-ascorbic acid (Wako, 013-12061), 50 ng/ml EGF (Peprotech, AF-100-15), 2 μM CHIR99021 (R&D #4423), 0.5 μM A83-01 (Peprotech, 90943360), and 1 μM SB431542 (BioVision). Cells were passaged with TrypLE every 3-4 days, and 50,000 cells were passaged to ensure sustained growth. All tests were carried out on hTSCs within 30 passages. When the cells in 2D hTSCs attained 80% confluency, they were cultured in hTSC media according to published protocol^43^. Following dissociation of individual cells using TrypLE, hTSCs were washed twice with Advanced DMEM/F12. At a final concentration of 72% Matrigel in Advanced DMEM/F12, 3,000 cells were suspended in 30 μL of Matrigel droplets. Each of the 24 well plates were used to plant droplets. Before polymerizing the Matrigel droplets at 37°C for 30 minutes, a two-minute incubation step was necessary on the tabletop. After taking the plates out of the incubator, 500 μL of trophoblast organoid media (TOM) was added. TOM medium was produced as described earlier^43^.

### Plasmids and transfection

SARS-CoV-2 expression plasmids were acquired from Addgene, courtesy of the contribution from Dr. Nevan J. Krogan at the University of California, San Francisco. These constructs are based on the pLVX-EF1alpha vector, fused with a 2xStrep-tag (#141395). The ORF3a-mCherry plasmid, featuring a CMV promoter within a pmCherryN1 vector backbone, was also obtained from Addgene (#165138). Trophoblast cells were transfected with plasmids encoding SARS-CoV-2 viral proteins using Lipofectamine 3000 (Thermo, L3000015) in accordance with the manufacturer’s instructions. All experiments were performed under biosafety level 2 (BSL2) conditions.

### Lentivirus preparation

In 10 cm dishes, HEK293T cells were seeded at a density that would result in 70–80% confluence the next day. TransLentiX (TransLentiX Inc.) was used to perform transfection in accordance with the manufacturer’s instructions. A DNA combination was generated in a 4:3:1 ratio, containing 10 μg of the lentiviral transfer plasmid (ORF3a-WT and ORF3a-NoPBM), 7.5 μg of the packaging plasmid psPAX2, and 2.5 μg of the envelope plasmid pMD2.G. After adding this mixture and TransLentiX reagent to the cells, they were cultured for six to eight hours. Following incubation, fresh DMEM containing 10% foetal bovine serum (FBS) was added to the transfected medium. After that, the cells were left to generate the virus for 48–72 hours. The lentiviral particle-containing supernatant was then collected, and any cellular debris was removed by filtering it through a 0.45 μm filter. The virus was concentrated using lentiviral concentration reagent from Takara (Lenti-X) as described by manufacturer protocol. Before being used again, the virus particles were separated and kept at -80°C.

### Immunofluorescence

Immunofluorescence staining was conducted on both cell cultures and 3D SC-TOs. For cell cultures, Jeg-3 cells in six well glass bottom plates (Cellvis, P06-1.5H-N) post-transfection were fixed with 4% paraformaldehyde at room temperature for 15 minutes, permeabilized with 0.2% Triton X-100 for 10 minutes and blocked with 1% Bovine serum albumin (BSA) for one hour at room temperature to prevent non-specific binding. e antigens. The following primary antibodies were used: GAPDH (Cell Signaling Technology, 97166S & 5174S), LC3 (Novus, NB600-1384), SQSTM1/p62 (Abcam, ab91526), Human Chorionic Gonadotropin (Abcam, ab9582), ZO1 (Proteintech, 21773-1-AP), Anti-strep tag (Sigma, SAB2702216), ORF3a (Cell Signaling Technology, 34340 & R&D Systems, MAB10706. Secondary antibodies used in the study were Alexa Fluor 488 goat anti-mouse IgG1 and goat anti-rabbit IgG (Thermo, A-11001 & A-11008), Alexa Fluor 594 goat anti-mouse and goat-anti-rabbit (Thermo, A-11005 & A-11012) and Alexa Fluor 647 goat anti-mouse and goat-anti-rabbit (Thermo, A21237 & A-21245).

Paraffin-embedded fixed placental villous tissue blocks were cut into 5-micrometer-thin sections which further underwent deparaffinization, rehydration, and antigen retrieval in a citrate buffer. The tissue sections were incubated with 1%BSA for blocking and subsequently treated with primary antibodies diluted in 1%BSA targeted to specific antigens for overnight at 4^0^C, followed by appropriate fluorescently conjugated secondary antibodies diluted in 1% BSA for 1 hour at room temperature. Nuclei were counterstained with Hoechst-33342 (Invitrogen, H3570) according to the manufacturer’s instructions. After the incubation, sections and cells were washed with PBS and mounted using Prolong Gold (Thermo, P36930) anti-fade mounting medium. The prepared samples were then examined under ECLIPSE Ti2 (Nikon) confocal microscope to elucidate the presence of respective antigens.

### Proteostat staining

Cultured human trophoblast cells were fixed using 4% paraformaldehyde for 15 minutes at room temperature for proteostat aggresome detection. Simultaneously, paraffin-embedded sections, rehydrated and deparaffinized, were prepared alongside. Both fixed cells and tissue sections were permeabilized with 0.2% Triton X-100 and blocked with 1%BSA to prevent non-specific staining. The Proteostat aggresome (Enzo life sciences Inc, ENZ-51035-0025) staining proceeded as per the manufacturer’s guidelines, with an emphasis on protecting the samples from light. Proteostat dye was added at a 1:2000 dilution for 30 min at room temperature. Post-staining, cells and sections were washed, counterstained with Hoechst-33342 for nuclear visualization, and mounted using Prolong gold anti-fade mounting media for fluorescence microscopy.

### Invasion assay

For the cell invasion assay, HTR-8 cells were employed to investigate the invasive capacity following transfection with SARS-CoV-2 protein-encoding plasmids. Prior to the assay, cells were cultured under standard conditions in six well plates and transfected for 24 hours with the plasmids of interest. Post-incubation, cells were detached using trypsin-EDTA solution (Gibco, 25300054). The cells were then carefully counted to obtain accurate cell numbers for seeding. Each trans-well insert with precoated Matrigel (Corning, 354480) was placed in 24 well plates according to the manufacturer’s protocol. The number of cells seeded into each insert was standardized to ensure uniformity across experiments. After seeding, the cells were allowed to invade through the Matrigel coating towards a chemotactic gradient provided by the medium containing 20% fetal bovine serum (FBS: Gibco, 16140071) in the lower chamber.

The invasion process was conducted under optimal cell culture conditions (37 °C, 5% CO2) for a duration of 24 hours to ensure significant invasion while preventing over confluency. At the end of the invasion period, non-invading cells were removed from the upper surface of the membranes using cotton swabs. Cells that had invaded through the Matrigel and reached the lower surface of the membrane were fixed, stained, and quantified. The fixed cells were usually stained with crystal violet (Sigma, C0775) , which allows for easy visualization and counting under a microscope.

### Western blot

Post-transfection, JEG-3 cells were lysed after a 24-hour period using RIPA lysis buffer (Thermo, 89901), supplemented with cocktails of protease and phosphatase inhibitors for protein preservation. Protein concentrations were determined via the Bicinchoninic Acid (BCA) method (Thermo, 23225). The lysates were then subjected to electrophoresis on 4%-20% gradient Mini-PROTEAN TGX precast gels (Bio-Rad Laboratories, 4561093) for resolution of the proteins. Subsequently, proteins were electroblotted onto Polyvinylidene Difluoride (PVDF) membranes (Sigma, 05317-10EA) at a constant 60 volts for two hours. Following the transfer, membranes were blocked to prevent non-specific binding and incubated with the relevant primary antibodies at 4°C overnight. The following day, membranes were thoroughly rinsed three times with 0.1% Tween-20 in Tris-buffered saline (TBST) before a 1-hour room temperature incubation with IR dye-linked secondary antibodies. After a final washing step to remove unbound antibodies, protein bands were visualized on a ChemiDoc imaging system (Bio-Rad Laboratories), using infrared detection wavelengths. Band intensities were measured with Image Lab software (Bio-Rad), and data were normalized against housekeeping proteins (β-actin/GAPDH) to ensure accuracy in protein quantification.

### Statistical Analyses

All statistical analyses were conducted using GraphPad Prism 9. Sample sizes were not predetermined using statistical methods. The Shapiro-Wilk test was employed to verify the normal distribution of continuous variables. Pairwise comparisons were evaluated for statistical significance using either the Student’s t-test or the Mann-Whitney test, depending on their suitability. For comparisons involving three or more groups, a one-way ANOVA was utilized, followed by Tukey’s test for identifying group differences. In cases of nonparametric data distribution, the Kruskal–Wallis test was applied, along with Dunn’s test for post-hoc analysis. Detailed descriptions of the statistics, tests used, and post-hoc tests for multiple comparisons are included in the legends of each figure and their source data.

## Acknowledgments

This work was supported in part by NIH/National Institute of Child Health and Human Development grant R01HD091218-04S1 (to IUM and JCK), NIH/NIGMS training grant, 5T32GM136554-03 (to BRJ) and a grant to Baylor College of Medicine from the Howard Hughes Medical Institute through the Gilliam Fellows Program to BRJ; NIH/NICHD grant, R01 HD110408, NIH/NIAID grant, R01AI141501, and a NIH/NIGMS Cobre grant, 1P20 GM121298 (to SS). DK is supported by the Early Career Award Program grant from Thrasher Research Fund. The work in the Theunissen lab was supported by an NIH Director’s New Innovator Award (DP2 GM137418) and grants from the Shipley Foundation Program for Innovation in Stem Cell Science and the Edward Mallinckrodt, Jr. Foundation. JCK and ED were also supported by NIH/NIDA R21DA057493-02; and NIH/NICHD 1R01HD113199-01.

## Author contributions

DK and IUM conceived the experimental plan; DK performed the majority of experiments assisted by RMK, BRJ, ERM, SJ. TWT and SS provided their expertise. JK and ED provided human placental tissue samples from the STOP-COVID-19 study. DK and IUM wrote the manuscript, and all authors approved the final draft.

## Materials availability

Further information and requests for resources and reagents should be directed to and will be fulfilled by the lead contact, Indira Mysorekar

---

**Supplementary Figure 1:**
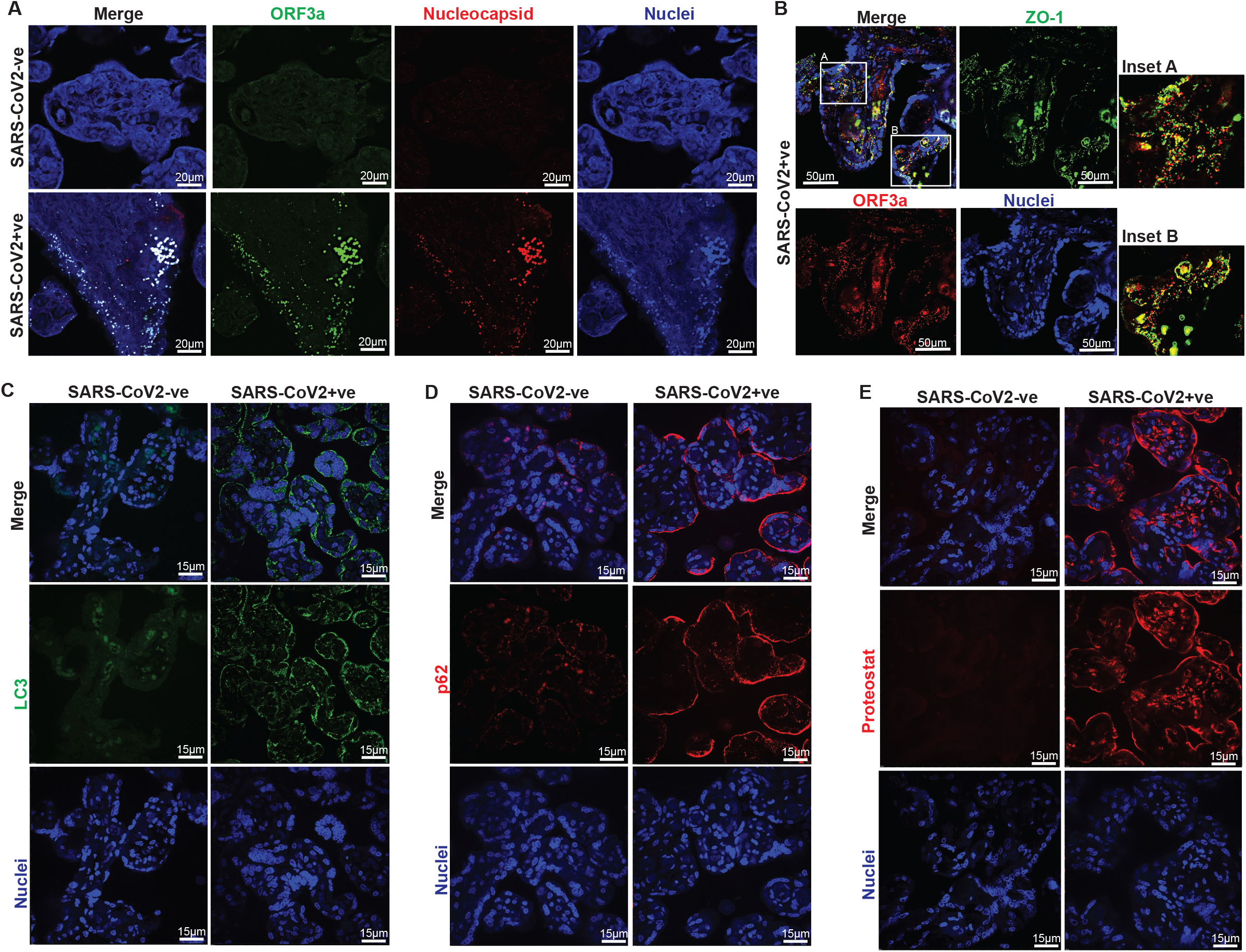
SARS-CoV-2 Infection of Human Term Placenta. **(A)** A representative confocal microscopy image illustrating the presence of ORF3a (green) and Nucleocapsid (red) in term placentas infected with SARS-CoV-2. Scale bar = 20 μm; 20X magnification objective**. (B)** A representative confocal microscopy image of SARS-CoV-2-infected term placenta demonstrates the co-localization of ORF3a (red) with the tight junction protein ZO-1 (green), producing a merged yellow signal (inset A and B). Scale bar = 50 μm; 40X magnification**. (C)** Immunostaining for the autophagy marker LC3 reveals heightened staining in SARS-CoV-2-infected term placenta relative to uninfected tissues. Scale bar = 15 μm; 60X magnification **(D)** Representative image illustrating increased immunostaining for the autophagy adaptor protein p62 in SARS-CoV-2-positive term placentas relative to uninfected controls. Scale bar = 15 μm; 60X magnification. **(E)** Representative confocal image illustrating enhanced Proteostat staining, signifying higher protein aggregation, in SARS-CoV-2-positive placentas relative to uninfected term placentas. Scale bar = 15 μm; 60X magnification.

